# Absolute measurement of the tissue origins of cell-free DNA in the healthy state and following paracetamol overdose

**DOI:** 10.1101/715888

**Authors:** Danny Laurent, Fiona Semple, Philip J. Starkey Lewis, Elaine Rose, Holly A. Black, Stuart J. Forbes, Mark J. Arends, James W. Dear, Timothy J. Aitman

## Abstract

**Background:** Despite the emergence of cell-free DNA (cfDNA) as a clinical biomarker in cancer, the tissue origins of cfDNA in healthy individuals have to date been inferred only by indirect and relative measurement methods, such as tissue-specific methylation and nucleosomal profiling.

**Methods:** We performed the first direct, absolute measurement of the tissue origins of cfDNA, using tissue-specific knockout mouse strains, in both healthy mice and following paracetamol (APAP) overdose. We then investigated the utility of total cfDNA and the percentage of liver-specific cfDNA as clinical biomarkers in patients presenting with APAP overdose.

**Results:** Analysis of cfDNA from healthy tissue-specific knockout mice showed that cfDNA originates predominantly from white and red blood cell lineages, with minor contribution from hepatocytes, and no detectable contribution from skeletal and cardiac muscle. Following APAP overdose in mice, total plasma cfDNA and the percentage fraction originating from hepatocytes increased by ~100 and ~19-fold respectively. Total cfDNA increased by an average of more than 236-fold in clinical samples from APAP overdose patients with biochemical evidence of liver injury, and 18-fold in patients without biochemically apparent liver injury. Measurement of liver-specific cfDNA, using droplet digital PCR and methylation analysis, revealed that the contribution of liver to cfDNA was increased by an average of 175-fold in APAP overdose patients with biochemically apparent liver injury compared to healthy subjects, but was not increased in overdose patients with normal liver function tests.

**Conclusions:** We present a novel method for measurement of the tissue origins of cfDNA in healthy and disease states and demonstrate the potential of cfDNA as a clinical biomarker in APAP overdose.

## Background

In 1948, Mandel and Metaìs described the presence of DNA in plasma [1], now known as cell-free DNA (cfDNA) and known to be present in other bodily fluids, such as urine [2], saliva and cerebrospinal fluid [3]. Release of cfDNA into the circulation is likely due to cellular breakdown mechanisms, such as apoptosis and necrosis, as well as active DNA release mechanisms [4]. The size profile of cfDNA typically follows multiples of the ~180bp nucleosomal unit[5, 6], with a dominant peak size of 167bp [7].

Analysis of cfDNA is clinically useful for detecting genetic and epigenetic alterations in DNA by allowing repeated non-invasive or minimally invasive sampling from bodily fluids [8]. Analysis of foetal cfDNA in maternal plasma has been clinically implemented for non-invasive prenatal testing of chromosomal abnormalities and some monogenic disorders [9, 10]. Analysis of cfDNA in cancer genomics is evolving rapidly [11–14] and utility of cfDNA as a biomarker of internal tissue damage [15–17], in organ transplantation [18] and in autoimmune disease is being investigated [19, 20].

Despite these developments, the tissue origins of cfDNA are not yet completely known. Current understanding rests on indirect and relative measurement methods, such as tissue-specific methylation markers [21–23] and nucleosomal profiling [7], with substantial variations in the estimated proportions of the tissues of origin between different studies [24]. Here, we present the first absolute, direct measurement of the tissue origins of cfDNA using six tissue-specific knockout mouse strains. We demonstrate that hepatocytes are a minor contributor to the pool of circulating cfDNA in healthy mice, and show a substantial increase in hepatocyte-derived cfDNA in mice exposed to an overdose of paracetamol (also known as acetaminophen, acetyl-para-aminophenol or APAP). Ultimately, we demonstrate the potential of cfDNA as a clinical biomarker in APAP overdose in human subjects. Overall, our findings form an important foundation for the development of cfDNA-based assays as biomarkers in non-malignant human disease.

## Methods

### Generation of tissue-specific knockout mice

Absolute measurement of the tissue origins of cfDNA in healthy mice was achieved by analysing cfDNA from tissue-specific knockout mice. cfDNA originating from the cells of interest, which express Cre recombinase from a cell-specific promoter, were recombined (1lox), as opposed to unrecombined (2lox) from other cell types in the body that do not express Cre recombinase (**Additional File 1: Figure S1**). For this study, a conditional mutation in the floxed *mT/mG* dual reporter gene [25] was generated in C57BL/6 mice by crossing mice homozygous for the floxed *mT/mG* gene with mice expressing different cell/tissue-specific Cre recombinase to obtain cell/tissue-specific knockout F1 mice. Myeloid (LysMCre) [26], lymphoid (hCD2-iCre) [27], cardiomyocyte (cTnTCre) [28], hepatocyte (AlbCre) [29] and striated muscle (MCKCre) [30] Cre mice were obtained from the Jackson laboratory. Erythroid (EpoRCre) [31] mice were obtained from Prof. Stuart Orkin (Cooperative Centers of Excellence in Hematology, Bostons Children’s Hospital). Mouse breeding was carried out in individually ventilated cages (4-6 mice per cage depending on litter size) with constant access to food and water, in the Biomedical Research Facility (BRF), University of Edinburgh (UoE) under Home Office project license PPL P1070AFA9 and genotyping was performed by Transnetyx (Cordova, Tennessee, USA) to confirm the presence of Cre recombinase and the floxed *mT/mG* gene. Male tissue-specific knockout mice (F1, n > 10 for each knockout line, Additional File 2: Figure S2) were culled at 10-12 weeks old via exposure to gradually increasing concentrations of carbon dioxide gas (schedule 1 method) at 10% displacement rate (chamber volume / minute). Carbon dioxide flow was continued for 1-2 minutes after cessation of breathing. No anaesthetic agent was used in the study. Blood was collected post-mortem from the inferior vena cava using 1ml syringes with a 25G needle into EDTA tubes. Plasma was obtained by centrifugation of blood at 1000g for 10 minutes, then 16000g for 5 minutes, within 3 hours of blood collection. Tissue samples were collected from mice post-mortem and stored at −70°C prior to analysis.

### Induction of APAP overdose in mice

Mice were fasted 12 hours prior to intraperitoneal (IP) injection with APAP (350 mg/kg mouse body weight), maintained in a 30°C incubator and given semi-solid food. Experiments were carried out under the Home Office project license PPL 70/7847. To investigate the timepoint where cfDNA concentration is highest following APAP overdose, culling and plasma collection were performed at 8, 24 and 48 hours following APAP injection in C57BL/6 mice (Charles River Laboratories, UK; n = 4 per time point). Contribution of hepatocytes to cfDNA levels was determined at 8 hours post APAP injection in hepatocyte-specific knockout mice (n = 4). Negative control mice injected with saline were included in each experiment. Mouse treatments were assigned prior to measurement of mouse body weight with restricted randomisation (each cage contained APAP and saline injected mice). Dosing for all mice was performed during the day within an hour. Mouse euthanasia was performed by exposure to increasing concentrations of carbon dioxide gas (as above). To confirm tissue damage from APAP dosing, liver tissues were sectioned and stained with hematoxylin and eosin (H&E) at the Pathology department, UoE, and liver function tests (alanine aminotransferase / ALT, aspartate aminotransferase / AST, albumin (ALB), bilirubin) were performed on plasma samples at the Shared University Research Facility, UoE.

### Clinical samples

Serum was collected from healthy volunteers (n = 11) and APAP overdose patients (n = 8) from the SNAP Clinical Trial [32]. The overdose group was comprised of two groups: normal ALT (n = 4) and raised ALT (n = 4), indicating clinical/biochemically apparent liver injury. Blood samples were centrifuged at 1000*g* for 15 minutes at 4°C and the supernatant was separated into aliquots and stored at −80°C prior to analysis. The protocol was approved by the Scotland A Research Ethics Committee, UK (ref no 10/MRE00/20).

### Genomic and cell-free DNA extraction and quantitation

Genomic DNA (gDNA) was extracted from frozen tissue samples using DNEasy blood and tissue kit (QIAGEN) and quantified using the Qubit dsDNA Broad Range assay kit (Thermo Fisher Scientific) according to manufacturer’s instructions. cfDNA was extracted from plasma using the QIAamp circulating nucleic acid kit (QIAGEN) according to the manufacturer’s instructions. cfDNA from clinical samples and mice was quantified using a single locus qPCR assay on the beta-actin (*ACTB* / *Actb*) gene [33, 34]. Fragment analysis of cfDNA was performed using Agilent DNA Bioanalyser.

### Droplet-digital PCR assay design and validation

A probe-based droplet-digital PCR (ddPCR) assay was designed to quantify the 1lox and 2lox *mT/mG* alleles. The assay was adapted from a probe-based qPCR assay [35] and designed using Primer3 [36] following the Biorad ddPCR application guide, with a maximum amplicon size of 130bp to accommodate for small fragments of cfDNA. The assay was validated by amplification of gDNA containing only 1lox or 2lox DNA fragments by ddPCR (**Additional File 3: Table S1**).

### Visualisation of Cre recombination with fluorescence imaging and histological analysis of tissue samples

Fluorescence microscopy of frozen tissue sections from knockout mice was performed to visually confirm Cre recombination in tissues, based on expression of cell membrane-localised enhanced green fluorescent protein (mG) and cell membrane-localised tdTomato (mT). Sections of 5μm were obtained from frozen tissues using a cryostat. These sections were put on a slide with mounting medium prolong gold and DAPI to stain nuclear DNA and left overnight at 4°C. Sectioning and staining were performed in the Pathology Department, UoE. Slides were visualised using a Britemac epifluorescence microscope in the Institute of Genetics and Molecular Medicine (IGMM) Advanced Imaging facility, UoE, with non-fluorescent frozen tissue slides from C57BL/6 mice as a negative control for tissue autofluorescence.

### Analysis of Cre recombination in genomic DNA and cell-free DNA

Cre recombination was analysed in gDNA of tissue-specific knockout mice using ddPCR to ensure successful recombination in the target tissue and check for specificity of Cre recombination in non-target tissues. Analysis of Cre recombination in the cfDNA of tissue-specific knockout mice was performed to show the contribution of the corresponding Cre-driven cell/tissue types to cfDNA. An input of 10ng of cfDNA was required for each well of ddPCR. Amplification of 1lox and 2lox alleles were performed in separate wells. Each sample was measured at least in triplicate. The amplification was run on a thermal cycler as follows: 10 min of activation at 95°C, 40 cycles of a two-step amplification protocol (30s at 94°C denaturation and 60s at 60°C), and a 10-min inactivation step at 98 °C. The metric used to show Cre recombination was percentage recombination (1lox%).

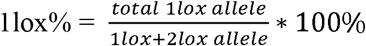

### Analysis of liver-specific methylation fragments in human cfDNA

The contribution of liver tissue to cfDNA in clinical samples was analysed using a methylation-specific ddPCR assay [37] on bisulphite-converted cfDNA using the Epitect DNA bisulphite conversion kit (QIAGEN).

### Statistics

Non-parametric statistical methods were performed on datasets to account for small sample sizes, the presence of data outliers, and data distribution that was not normal. Normality of data distribution was confirmed using Shapiro-Wilk test. The presence of outliers in datasets were confirmed using graphical examination of boxplots. A Mann-Whitney U test were performed to compare biomarker measurements between different groups of clinical samples, and to compare percentage of recombination of the floxed *mT/mG* gene in mouse tissues. Statistical dependence between biomarkers were assessed using Spearman’s rank correlation coefficient.

## Results

### Validation of ddPCR assay for absolute quantification of cfDNA tissue origins

We designed and validated a ddPCR assay to quantify recombination in the floxed *mT/mG* gene. Amplification of gDNA was performed to demonstrate specificity for each target allele (**Fig. 1a** **and** **b**). Amplification of 2 lox gDNA with the 1lox assay showed no positive droplet (**Fig. 1a**). Amplification of the 1lox gDNA with the 2lox assay showed 13 positive droplets out of a total of 16467 (less than 0.1%) (**Fig. 1b**). Assay sensitivity was tested using a dilution series of target alleles to show that the assay reproducibly quantifies less than 30 target alleles (**Fig. 1c**). Non-specific amplification was not observed in C57BL/6 background gDNA, nor in the water negative control.

**Fig. 1.**
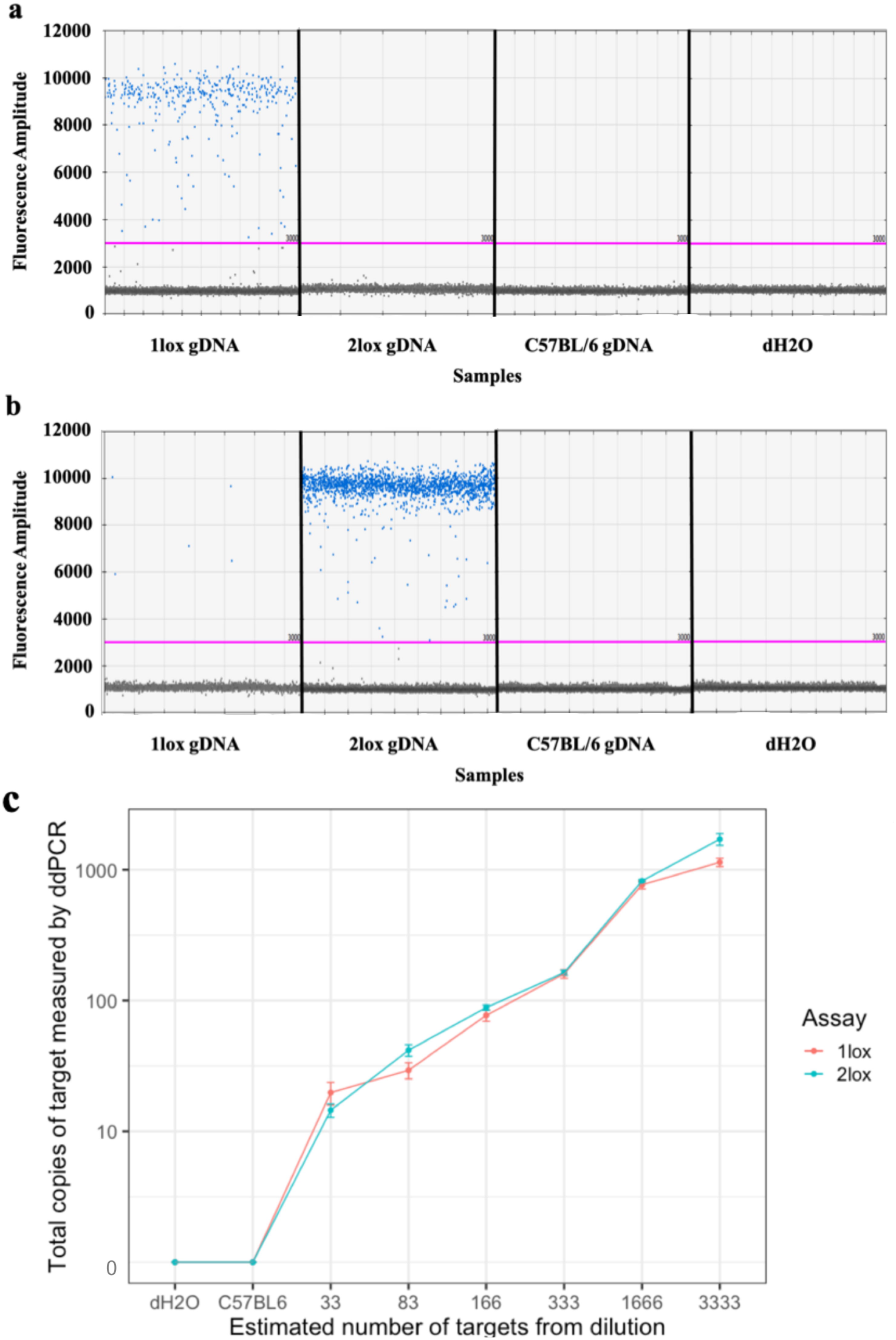
Validation of ddPCR assays on gDNA containing 1lox or 2lox alleles. (**a**) 1lox ddPCR assay specifically amplified gDNA containing 1lox alleles, (**b**) 2lox ddPCR assay amplified gDNA containing 2lox alleles with less than 0.1% of positive droplets from 1lox gDNA. The pink line shows the threshold between positive and negative droplets at fluorescence amplitude 3000. (**c**) Dilution experiment showed both assays robustly detected less than 30 targets. Each dot showed mean of 1lox or 2lox copies measured by ddPCR assays in six replicates. Error bars showed standard error.

### Analysis of DNA recombination in mouse tissues

Cre recombination was confirmed in expected tissues using ddPCR (**Fig. 2a**) and fluorescence microscopy based on the expression of the *mT/mG* reporter gene (**Fig. 2b**). To check for specificity of Cre recombination, we performed ddPCR on gDNA extracted from 16 tissue types for each knockout line. Highest specificity was found in hepatocyte and cardiomyocyte knockout lines, where recombination was observed in liver and heart, respectively, with minimal recombination in other tissues (**Fig. 2c**). Specificity of Cre recombination was more variable, but generally lower in the other knockout lines (**Additional File 4: Figure S3**).

**Fig. 2.**
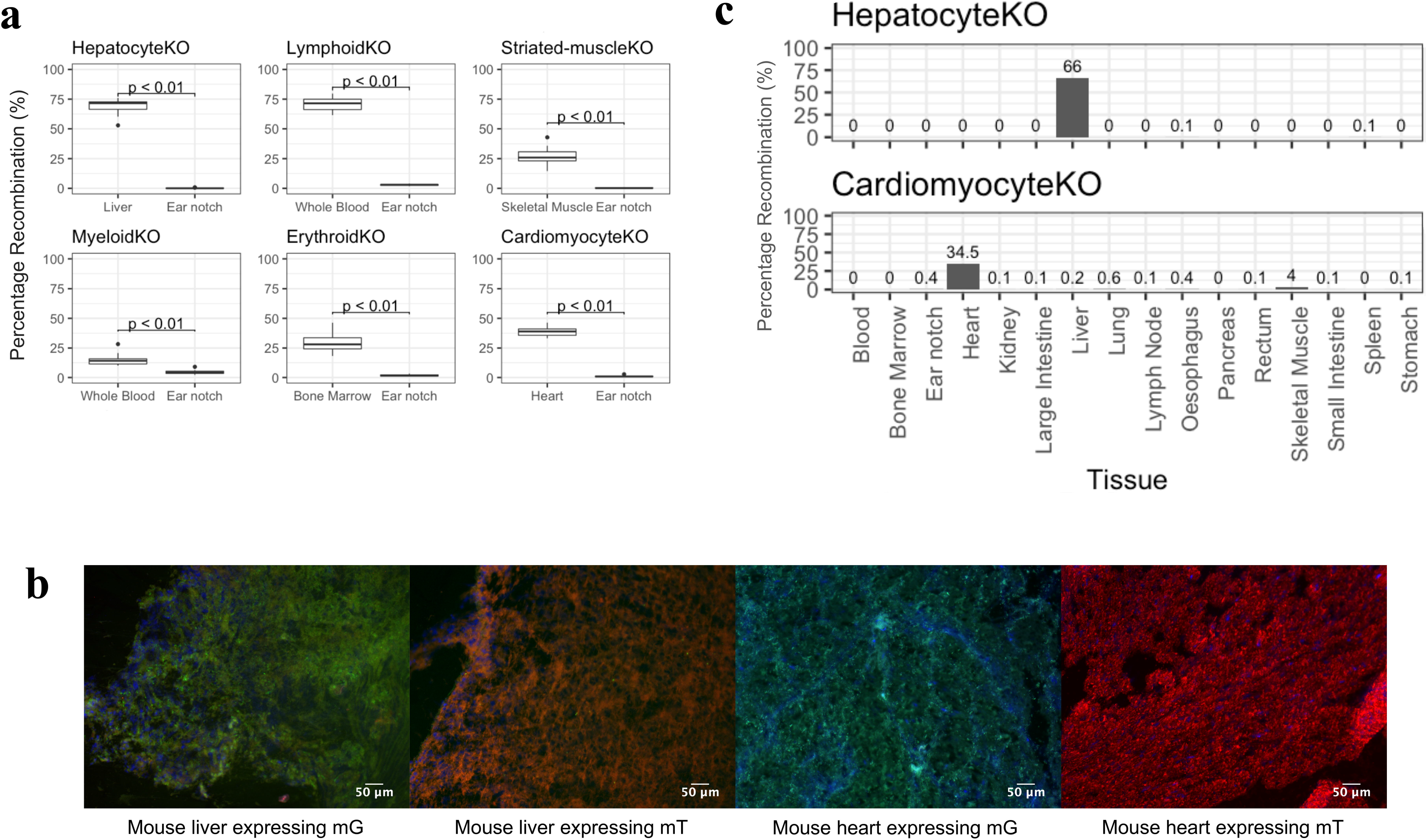
Analysis of Cre recombination in tissues of knockout mouse lines. (**a**) ddPCR measurement of 1lox allele and 2lox allele in tissues where recombination was expected vs. not expected for each knockout lines (n > 10 mice for each line). p-value obtained from a Mann-Whitney U test (**b**) Representative images for confirmation of Cre recombination in tissues shown by fluorescence imaging. Left to right: liver of a *AlbCre*;*mT/mG* mouse, liver of a *mT/mG* mouse, heart of a *cTnTCre*;*mT/mG* mouse, heart of a *mT/mG* mouse. Green colour in the tissues derived from the expression of mG, and red from mT. (**c**) Representative bar charts of % recombination showing specificity of Cre recombination in a hepatocyte- and cardiomyocyte-specific knockout mouse across many tissues shown by ddPCR assay.

### Tissue origins of cfDNA in healthy mice

To determine the tissue origins of cfDNA, we analysed cfDNA from the six tissue-specific knockout lines using ddPCR. Plasma from 10 or more mice for each knockout line was pooled prior to cfDNA extraction, to account for the small circulating blood volume of individual mice and the low physiological concentration of cfDNA (**Additional File 5: Figure S4**). Sufficient DNA was extracted for triplicate measurements of the pooled sample from each line by ddPCR (**Fig. 3a**). DNA fragment analysis confirmed the expected cfDNA fragment size profile which followed the mono-, di- and tri-nucleosome pattern associated with apoptosis, with larger cfDNA fragments most likely due to tissue necrosis and lysis (**Fig. 3b**).

**Fig. 3.**
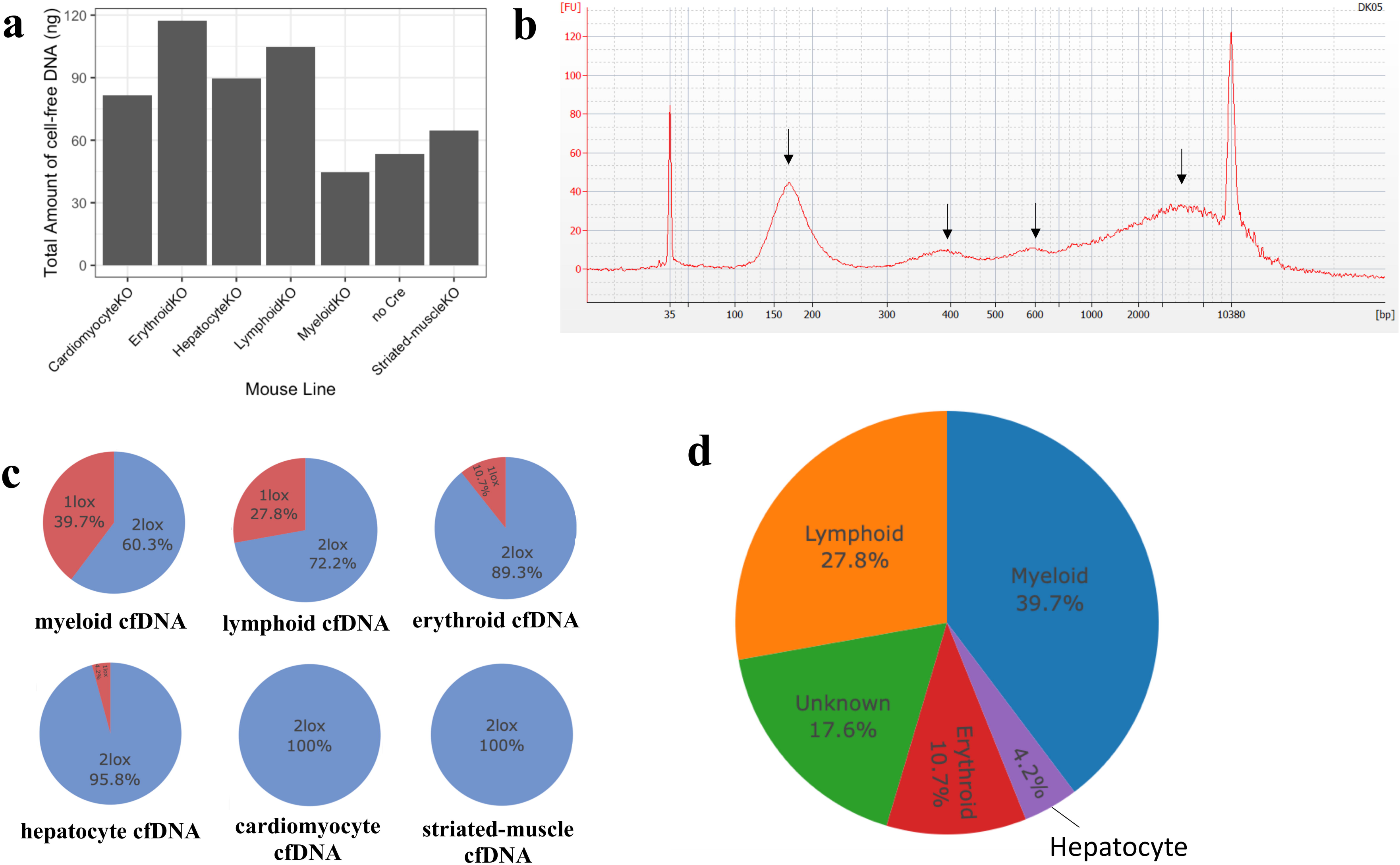
The tissue origins of cfDNA in healthy mouse models. (**a)**Total amount of cfDNA obtained from a minimum of 10 mice (pooled) per knockout line measured by qPCR of *ACTB* gene. (**b**) Representative cfDNA fragment analysis from hepatocyte-specific knockout mice. Fragments of cfDNA followed mono-, di-, and tri- nucleosome fragment sizes, and large cfDNA fragment size, shown by black arrows (**c**) Percentage recombination (1lox%) for each knockout mouse line showing contribution of different cell types to total cfDNA. (**d**) A cumulative percentage of cell/tissue contributions to cfDNA from ddPCR.

Contribution of different cell types to cfDNA was quantified by counting the number of 1lox and 2lox alleles in the cfDNA of mice for each knockout line and subsequently the percentage recombination (1lox%) for each of the knockout lines was calculated (**Fig. 3c**). Absolute measurement of cfDNA showed that myeloid, lymphoid and erythroid lineages were the major contributors to the levels of cfDNA in healthy mice, contributing 39.7%, 27.8% and 10.7% respectively, with a small contribution from hepatocytes (4.2%). Both cardiomyocyte and striated muscle cells had undetectable contribution to the levels of cfDNA (**Fig. 3d**).

### Tissue origins of cfDNA following tissue injury in mice

To validate tissue-specific knockout mice as a model system for studying the tissue origins of cfDNA, we measured hepatocyte-specific cfDNA levels after APAP overdose in hepatocyte-specific knockout mice. The optimal timepoint for sample collection was determined using C57BL/6 mice after APAP administration. Following APAP administration, mice developed phenotypic signs of acute liver injury, characterised by hunched posture, with no severe signs of liver injury, including loss of mobility or abnormal respiration. We assayed plasma for protein biomarkers of liver function and analysed histological liver sections at 8, 24 and 48 hours after administration of APAP which confirmed liver injury from 8 hours after APAP dosing (**Additional File 6: Figure S5**). Analysis of total plasma cfDNA demonstrated an increase of ~100-fold in APAP-dosed mice compared to mice receiving saline injection, which peaked at 8 hours post APAP dose. cfDNA in hepatocyte-specific knockout mice was subsequently measured at 8 hours, where we also observed greater than 100-fold increase in the total cfDNA following APAP dosing, compared to control mice receiving saline only injections. An increase of the liver biomarkers, ALT and AST, and histological analysis of mouse liver tissues confirmed the presence of liver injury (**Fig. 4**). Furthermore, quantification of liver-specific cfDNA at 8 hours post APAP dosing in hepatocyte-specific knockout mice shows hepatocyte contribution to cfDNA increased to 77.9%, compared to 4.2% in non-treated healthy mice (Fig.4g, **Additional File 7: Figure S6**).

**Fig. 4.**
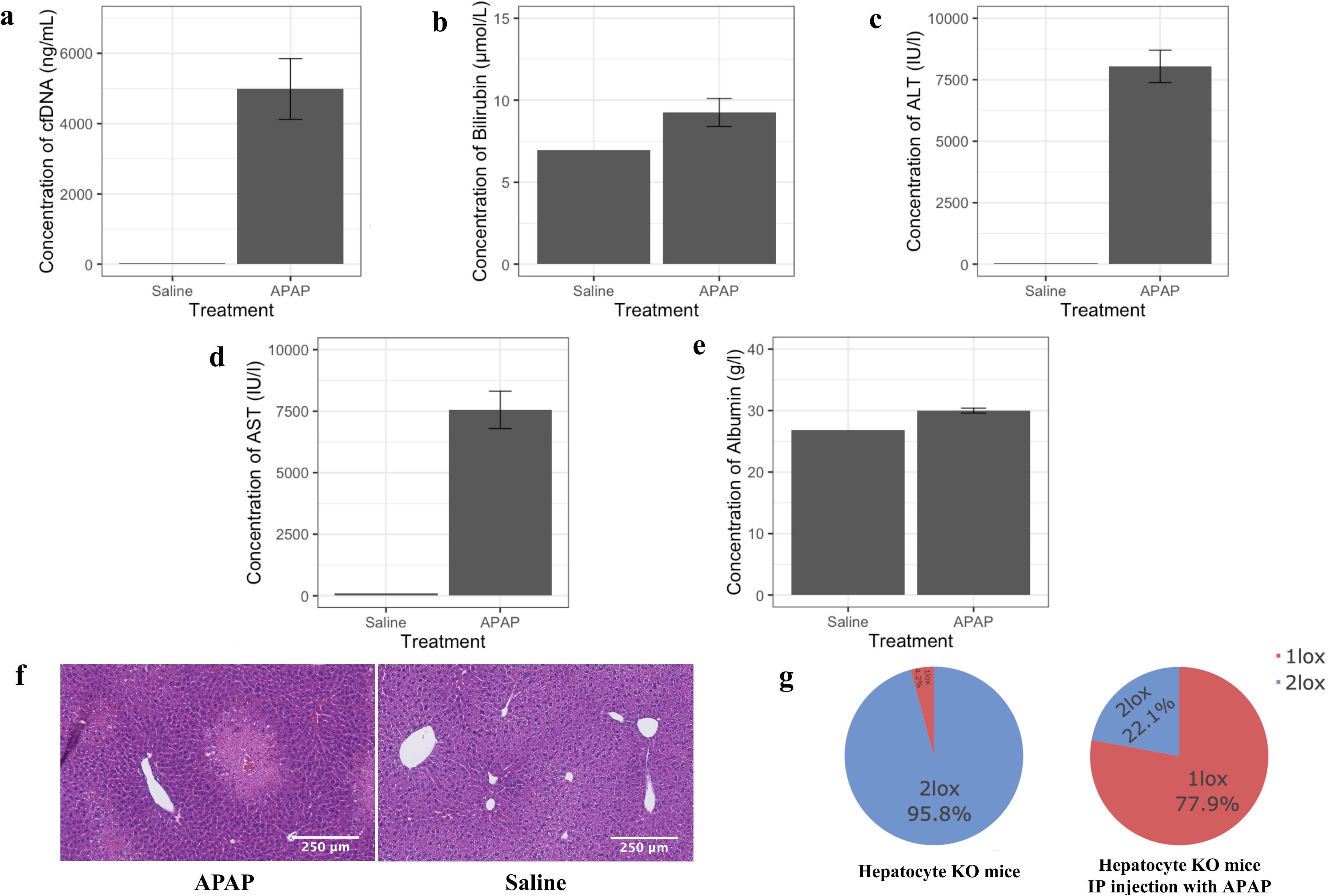
The tissue origins of cfDNA following APAP overdose in mouse models. Barplots of concentration of blood biomarkers from APAP-injected hepatocyte-specific knockout mice (n = 4) and saline-treated negative control (n = 10, pooled): (**a**) total cell-free DNA, (**b**) bilirubin, (**c**) AST, (**d**) ALT, (**e**) albumin. Data are shown as mean ± standard error. (**f**) H&E staining of liver tissues from an APAP-injected hepatocyte-specific knockout mice vs saline-treated negative control showing acute zone 3 coagulative necrosis typical of APAP hepatotoxicity. (**g**) Comparison of hepatocyte cell contribution in healthy mice vs post APAP overdose (n = 4).

### Analysis cfDNA in patients with APAP overdose

To demonstrate potential applicability of our findings to APAP overdose in humans, we measured the total amount of cfDNA and protein liver function biomarkers in the serum of healthy volunteers (n = 11) and APAP overdose patients (n=8), a subset of which (n = 4) were not exhibiting clinically apparent liver injury based on serum ALT levels (**Additional File 8: Table S2**). On average, the concentration of cfDNA in the APAP overdose patients increased by ~126-fold (**Fig. 5a**). In patients with clinically apparent liver injury (high ALT), total cfDNA increased by ~234-fold, whereas patients without clinically apparent liver injury (normal ALT) demonstrated mean total cfDNA increase of ~18-fold compared to healthy volunteers (**Fig. 5a**). The increase in total cfDNA following APAP overdose in patients was consistent with the increase in cfDNA following APAP in C57BL/6 and hepatocyte-specific knockout mice, indicating the potential of analysis of cfDNA as a clinical biomarker in APAP overdose.

**Fig. 5.**
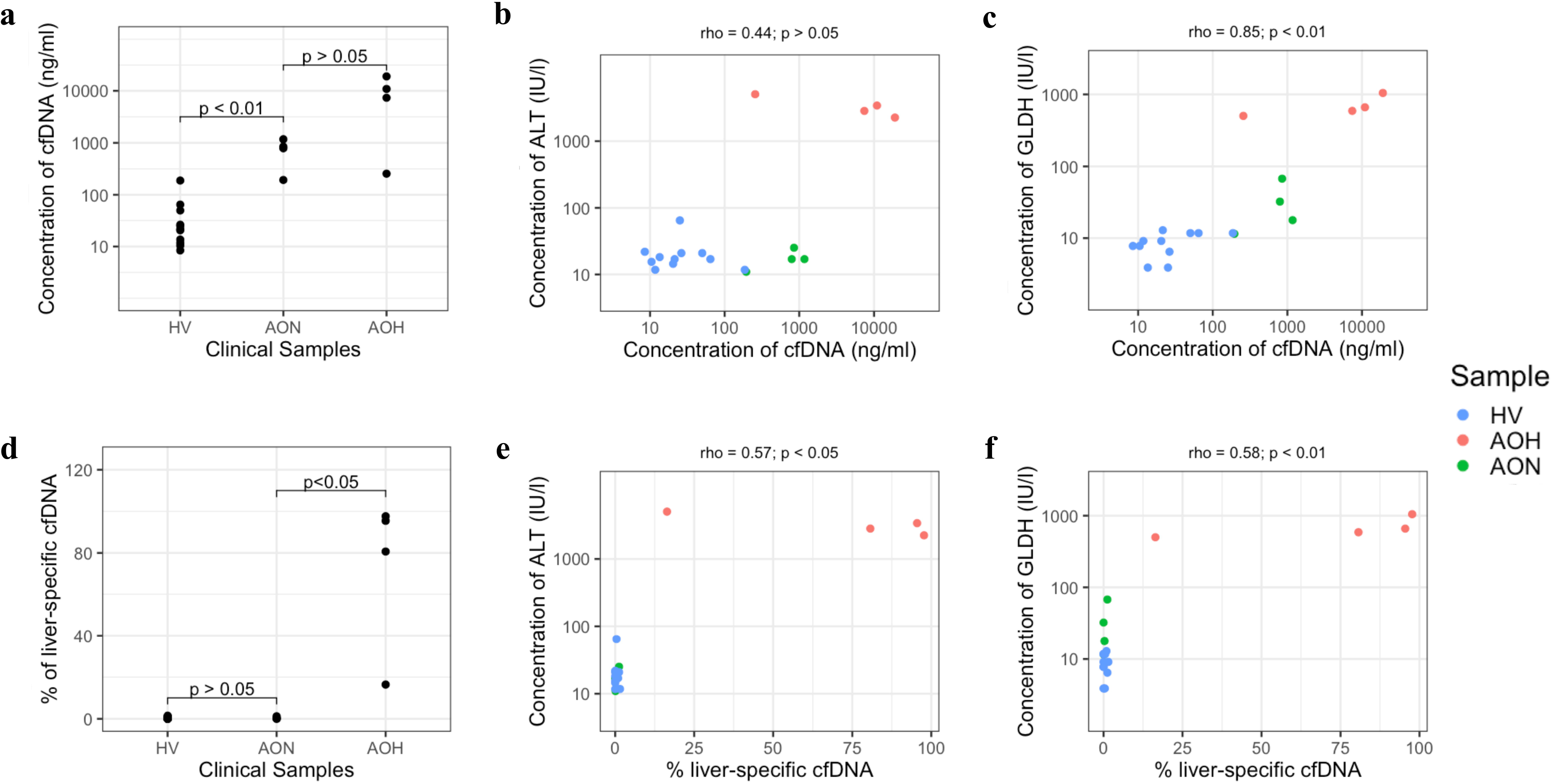
Analyses of cfDNA and other liver biomarkers in clinical samples. Comparison of (**a**) total concentration of cfDNA between healthy volunteers (HV), APAP overdose patients with normal ALT (AON) and high ALT (AOH) and (**b**) comparison of serum ALT and total concentrations of cfDNA, (**c**) comparison of serum GLDH and total concentrations of cfDNA, (**d**) comparison of percentage of liver-specific cfDNA between samples, (**e**) comparison of serum ALT and percentage of liver-specific cfDNA, (**f**) comparison of serum GLDH and percentage of liver-specific cfDNA. Statistical significance for panel (**a**) and (**d**) were obtained from Mann-Whitney U test and for panel (**b**), (**c**), (**e**), (**f**) from Spearman’s rank correlation coefficient.

To further assess the utility of cfDNA as a clinical biomarker, we compared the serum cfDNA concentration with other conventional biomarkers of liver function. Total cfDNA was higher in all patients with APAP overdose than in healthy volunteers, although two samples, one from an overdose patient with normal ALT (cfDNA = 192.87) and one healthy volunteer (cfDNA = 188.17) showed similar cfDNA concentrations (**Fig. 5b**). Total concentration of cfDNA also correlated with levels of glutamate dehydrogenase (GLDH), a novel APAP overdose biomarker that measures mitochondrial damage (Spearman correlation 0.85, p-value <0.01) with the (**Fig. 5c**).

Specific analysis of liver contributions to the cfDNA pool, measured by ddPCR, showed the percentage of liver-specific cfDNA fragments increased by ~175-fold in patients with clinically apparent liver injury compared to healthy volunteers, but no increase in liver-specific cfDNA was observed in overdose patients without clinically apparent liver injury (normal ALT) (**Fig. 5d, e**). Although GLDH levels were increased in APAP overdose patient with normal ALT, analysis of liver-specific cfDNA indicated that the cfDNA did not originate from liver tissue (**Fig. 5f**).

## Discussion

Our study demonstrates the utility of tissue-specific knockout mouse models as a platform to examine the tissue origins of cfDNA. Here, we present the first absolute, direct measurement of the tissue origins of cfDNA in healthy mice, as well as post-APAP overdose. Our data show that hematopoietic cells, including myeloid, lymphoid and erythroid cells, are the major components of cfDNA in the healthy state, with a minor contribution from liver and no detectable contribution from cardiomyocytes and skeletal muscle cells. Our results show concordance with other indirect methods, such as nucleosomal profiling and methylation analysis, that have assessed the tissue origins of cfDNA [7, 21–23], although these studies show considerable variation in the estimates of tissue origin, which may in part be due to the indirect or relative measurement methods [24]. Measurement of tissue origins of cfDNA in knockout mouse models is direct and specific for the tissue of interest and benefits from the ability to control external variables, further reducing sample variability. This experimental design may therefore be more accurate and less error-prone than indirect relative methods. We also demonstrate the applicability of this model system in a pathological scenario. The total concentration of cfDNA and the hepatocyte-derived fraction of cfDNA increased substantially after APAP-induced liver injury in mice. This large increase in total cfDNA and hepatocyte-specific cfDNA after APAP overdose demonstrate the sensitivity of the model system for detecting perturbations in the origins of cfDNA arising from external stimuli. The study is the first to show the contribution of a specific tissue to the pool of cfDNA using tissue-specific knockout mice, thereby serving as a generic proof of concept for investigations of the cellular origins of cfDNA in other physiological and pathological conditions. This may include exercise, infection, autoimmunity, trauma, myocardial infarction and transplantation as well as in studies of development and ageing. Analysis of the tissue origins of cfDNA has the potential to be applied to any condition associated with cell death or loss of nuclear DNA, into plasma or other bodily fluids, such as urine, saliva or cerebrospinal fluid.

APAP overdose is the most common cause of acute liver failure in the USA and Europe [38]. Previous studies have suggested the utility of cfDNA as a clinical biomarker of cell death in a range of clinical conditions [15, 18], but cfDNA has not previously been studied as a biomarker of liver damage in APAP overdose. We found that total concentration of cfDNA robustly distinguished APAP overdose patients from healthy volunteers, suggesting that total cfDNA concentration may reflect exposure to APAP. In overdose patients with raised ALT, the increase in cfDNA concentration was greater than in those with normal ALT, although all overdose patients had a higher cfDNA concentration than healthy volunteers. However, whilst there was also a clear increase in the proportion of cfDNA that originated from liver in overdose patients with raised ALT concentrations, the proportion of liver-specific cfDNA was not increased in overdose patients with normal ALT. This contrasts with our observation of increased GLDH in all overdose patients, both those with raised and normal ALT. Previous studies have proposed GLDH as a marker of liver damage in APAP overdose [39], but these data suggest that in APAP patients with normal ALT, the rise in total cfDNA and GLDH was not liver-derived and may have been due to damage of tissues other than the liver. Given that serum GLDH is a measure of mitochondrial damage [40] and that GLDH is expressed heterogeneously across mammalian tissues [41], the increase in GLDH and cfDNA in patients without clinically overt liver injury may reflect release of cfDNA from other tissues, for which further pre-clinical and clinical work is indicated to establish the source. Despite the novel approach of absolute and direct measurement of cfDNA, our study is limited by issues that are widely acknowledged in the field of cfDNA research [43]. Firstly, low cfDNA concentration and low plasma volume of mice means pooling of cfDNA from at least 10 mice was necessary to obtain sufficient material for ddPCR assay measurement. In addition, haemolysis and contamination of large cfDNA fragments that arises from lysis of blood cells during sample processing may have led to an overestimation of the fraction of cfDNA derived from white blood cells. Finally, the contribution of cells measured in this study is dependent upon the activity and specificity of Cre-recombinase, which varies between knockout lines.

Prior to wider application of cfDNA assays in the clinic, it will be important to understand the sources of variability in cfDNA concentrations in healthy subjects and the origins of cfDNA in the presence of physiological stimuli such as exercise and common disorders (e.g. upper respiratory infection). Whilst studies in humans can give indirect measures of tissue origins of cfDNA, optimal assay design and interpretation will require greater understanding of the effects of developmental stage and physiological and pathological state, as well as the mechanism of release for injured or dying tissue. These factors are difficult to study in humans and the model system reported here may yield insights that are of value for clinical assay development. For the specific clinical scenario studied here, APAP overdose, in which current patient stratification is acknowledged to be sub-optimal [44], follow-up studies in larger numbers will be needed for validation and formal assessment of sensitivity, specificity and predictive power.

## Conclusions

Absolute measurement of the tissue origins of cfDNA in the healthy state showed major contributions from hematopoietic cells, minor contributions from hepatocytes, and no detectable contribution from cardiac and skeletal muscle. The analysis of cfDNA is a candidate biomarker for stratification of APAP overdose patients.

## Abbreviations

ACTB: beta-actin
ALB: albumin
ALT: alanine aminotransferase
APAP: paracetamol/acetaminophen/N-acetyl-para-aminophenol
AST: aspartate aminotransferase
BHQ: black hole quencher
cfDNA: cell-free DNA
ddPCR: droplet-digital PCR
FAM: 6-carboxyfluorescein
gDNA: genomic DNA
GFP: green fluorescent protein
GLDH: glutamate dehydrogenase
H&E: hematoxylin and eosin
IP: intraperitoneal
mG: cell membrane-localised enhanced green fluorescent protein
mT: cell membrane-localised tdTomato
1lox: recombined floxed allele
1lox%: percentage recombination of floxed alleles
2lox: unrecombined floxed allele

## Declarations

### Ethics approval and consent to participate

Animal model work in this study was carried out in line with UK Home Office project licenses PPL P1070AFA9 and PPL 70/7847. The protocol for clinical samples was approved by the Scotland A Research Ethics Committee, UK (ref no 10/MRE00/20)

### Consent for publication

Not applicable

### Availability of data and materials

All data supporting the conclusion of this article are included in this published article and its additional files.

### Authors’ contributions

DL conceived and designed the study, performed experiments and analyses, and wrote the manuscript. FS and PJSL designed the study and performed experiments and analyses. ER performed experiments and analyses. HAB designed experiments and performed analyses. MJA performed analyses. SJF and JWD designed the study. TJA conceived and designed the study, performed analyses, wrote the manuscript and provided funding. All authors read and approved the final manuscript.

## Acknowledgements

We thank Prof. Mary Porteous and the South East Scotland Genetics service for access to the Biorad droplet-digital PCR system, Prof. Stuart Orkin of Cooperative Centers of Excellence in Hematology (CCEH) core laboratory (supported by National Institute of Diabetes and Digestive and Kidney Diseases), Boston Children’s Hospital, for his generous donation of EpoR Cre mice, and members of our laboratory for critical review of the manuscript.

## Additional Files

Additional File 1

File format: .jpg

Title of data: Figure S1. A schematic representation of the overall strategy for absolute quantitation of the tissue origins of cell-free DNA.

Description of data: A conditional knockout is generated in the tissue of interest (e.g. liver) for each mouse model, containing a tissue/cell-specific Cre recombinase and the reporter gene floxed *mT/mG*. Cre recombination occurs in target cells/tissues, causing deletion of a loxP site and the *mT* gene, leaving one loxP site in the DNA (1lox), and enabling the expression of the *mG* gene as opposed to other tissues in the body that contain two loxP sites (2lox). Histology and DNA analysis are performed in mouse tissues to confirm Cre recombination. Cell-free DNA, containing a mixture of 1lox and 2lox alleles released by various tissues through normal tissue turnover, can be extracted and analysed to reveal absolute contribution of cell-type/tissue of interest.

Additional File 2

File format: .tiff

Title of data: Figure S2. Body weight measurements from different mouse lines at 10-12 week old.

Description of data: Body weight of mice (n > 10) ranges between 20.6 and 31.9 gram. Tissue knockout mouse lines showed similar body weight to non-knockout control (no Cre).

Additional File 3

File format: .xls

Title of data: Table S1. Oligonucleotides for amplification of 1lox and 2lox *mT/mG* gene. Description of data: 1lox assay consisted of mTmGF1, mTmGR and mTmGP. 2lox assay consisted of mTmGF2, mTmGR, mTmGP. FAM: 6-carboxyfluorescein; BHQ: black hole quencher.

Additional File 4

File format: .tiff

Title of data: Figure S3. Specificity of Cre recombination across 16 mouse tissues in four knockout lines.

Description of data: Knockout lines from top to bottom: myeloid, lymphoid, erythroid and striated muscle.

Additional File 5

File format: .tiff

Title of data: Figure S4. Total amount of cfDNA extracted from a single mouse, and pools of 5, and 10 C57BL/6 mice.

Description of data: Total amount of cfDNA from a pool of 10 mice provided sufficient cfDNA input for at least duplicate measurements using ddPCR (20ng for a single measurement of 1lox and 2lox alleles).

Additional File 6

File format: .tiff

Title of data: Figure S5. The effect of APAP on biomarkers at different timepoints in C57BL/6 mice.

Description of data: The concentration of cfDNA (**A**), concentration of alanine aminotransferase (**B**) and aspartate aminotransferase (**C**) peaked at 8 hours following APAP dosing and decreased after 24 and 48 hours. The levels of albumin (**D**) and bilirubin (**E**) were unaffected by APAP dosing, indicating the absence of overt liver failure. Histological analysis (**F**) showed that liver damage (acute zone 3 coagulative necrosis typical of APAP hepatotoxicity) was most severe at 24-hour following APAP compared to other time points.

Additional File 7

File format: .tiff

Title of data: Figure S6. The effect of intraperitoneal injection on the tissue origins of cfDNA Description of data: The increase of hepatocyte contribution in the analysis of the tissue origins of cfDNA 8-hour after APAP dosing in hepatocyte-specific knockout mice was not caused by intraperitoneal injection, as shown by a decrease of tissue contribution in mice injected with saline.

Additional File 8

File format: .xls

Title of data: Table S2. Measurements of serum biomarkers from APAP overdose patients and healthy volunteers.

Description of data: Normal range for serum biomarker [45, 46]: Bilirubin (2-17 μmol/l), ALT (0-45 IU/l), AST (0-35 IU/l), ALP (30-120 IU/l), ALB/Albumin (40-60 g/l), GLDH (1-10 IU/l).

