## Supplementary figures and images for "Absolute measurement of the tissue origins of cell-free DNA in the healthy state and following paracetamol overdose"

### Additional File 1

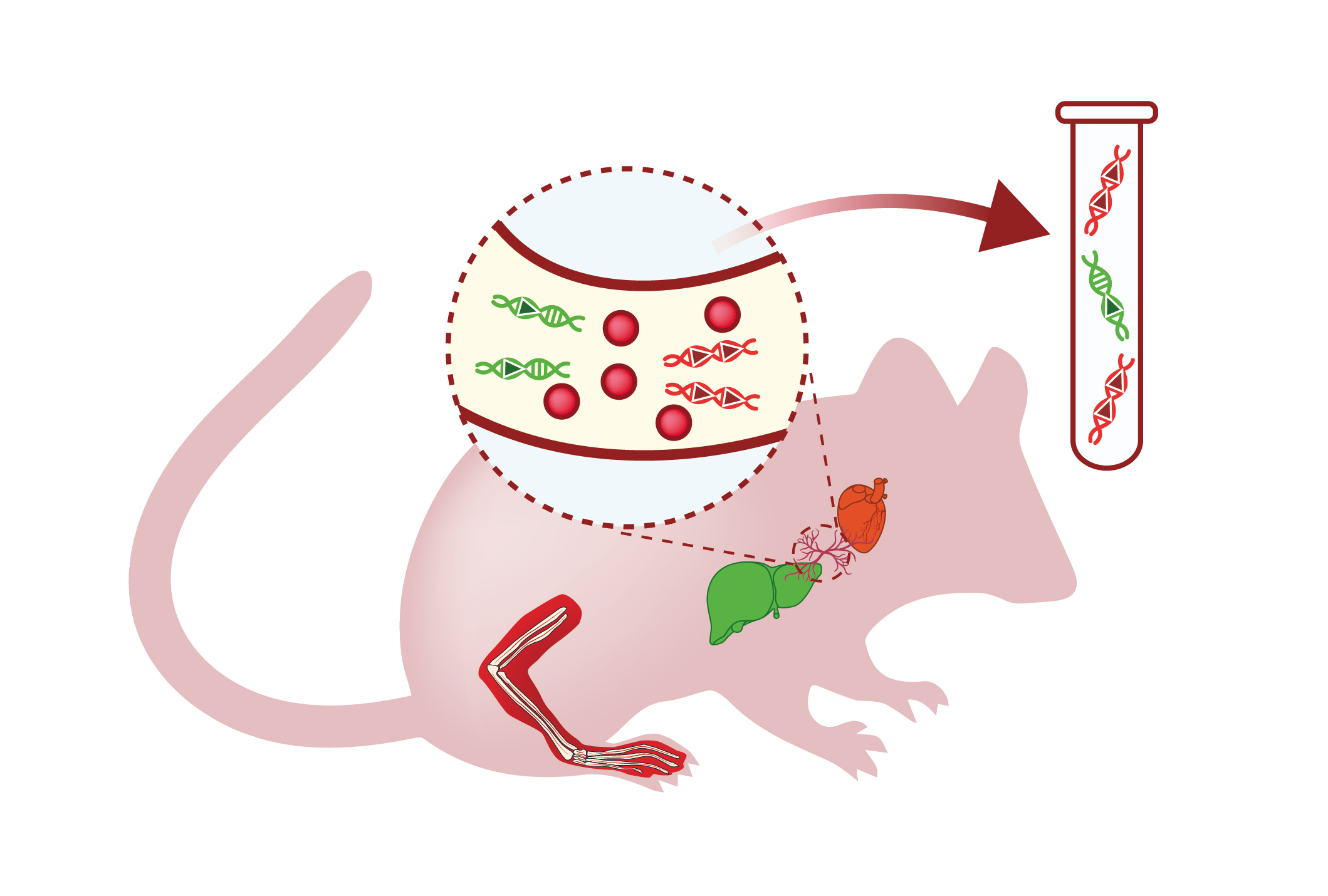

### Additional File 2

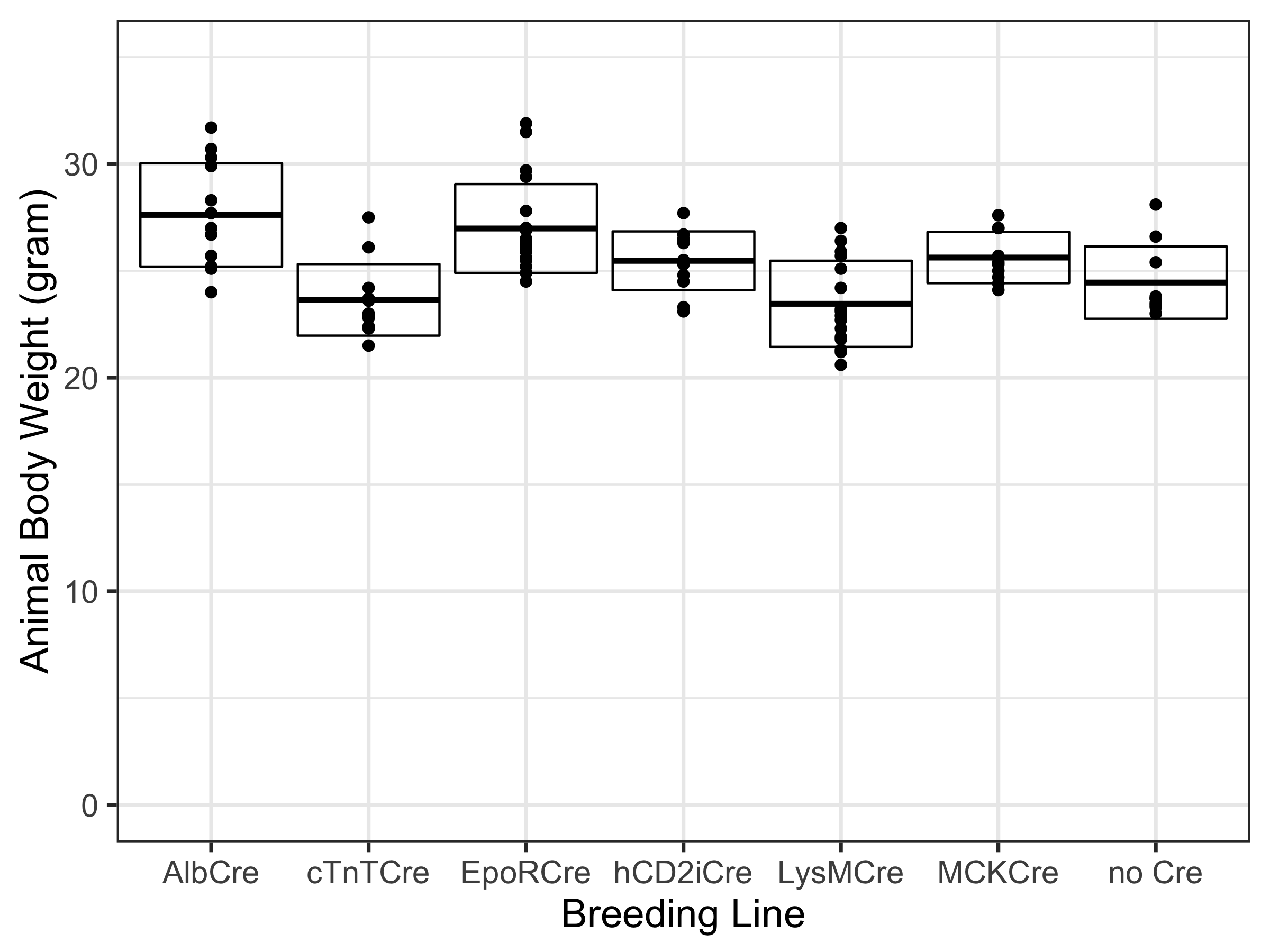

### Additional File 4

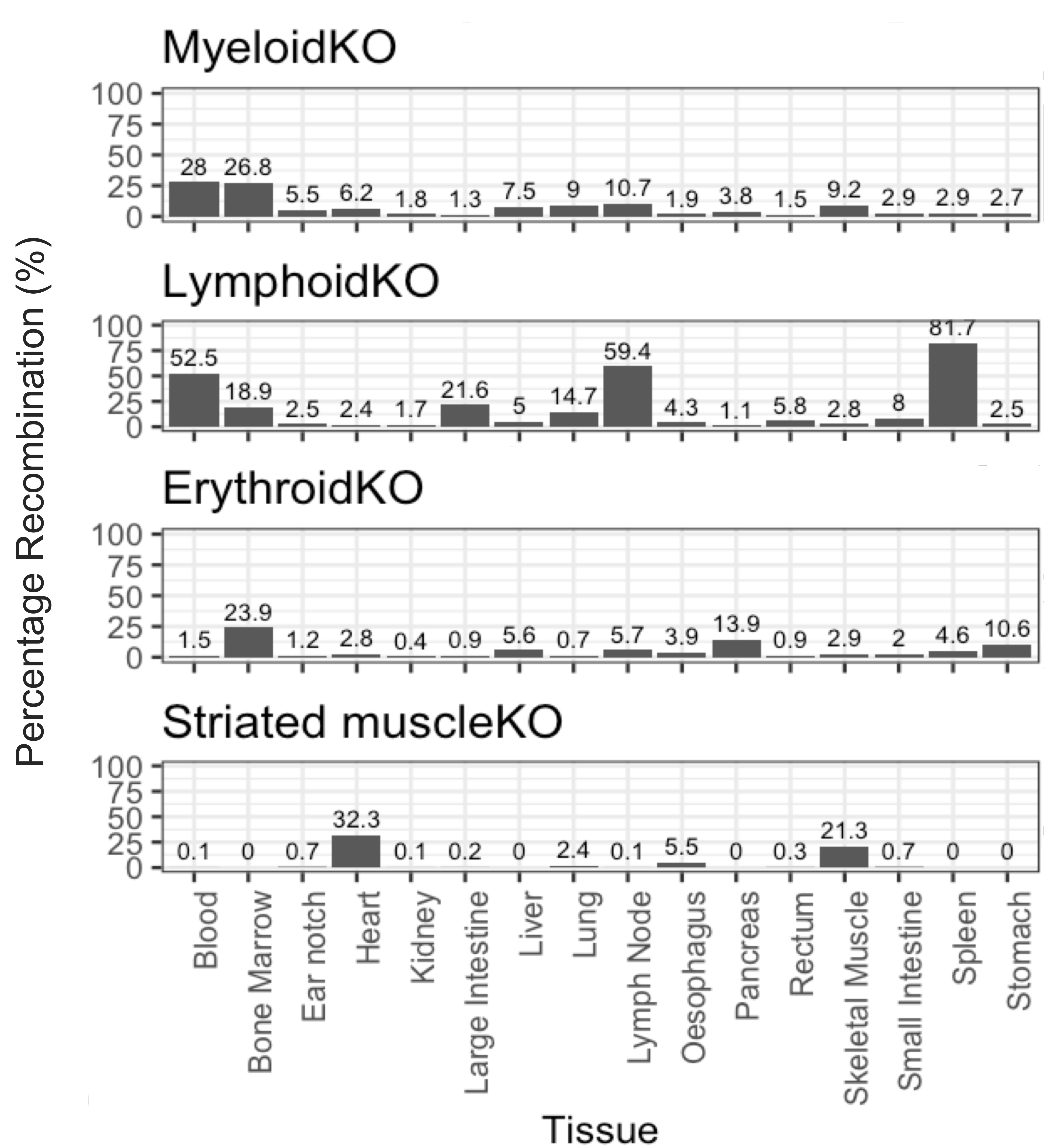

### Additional File 5

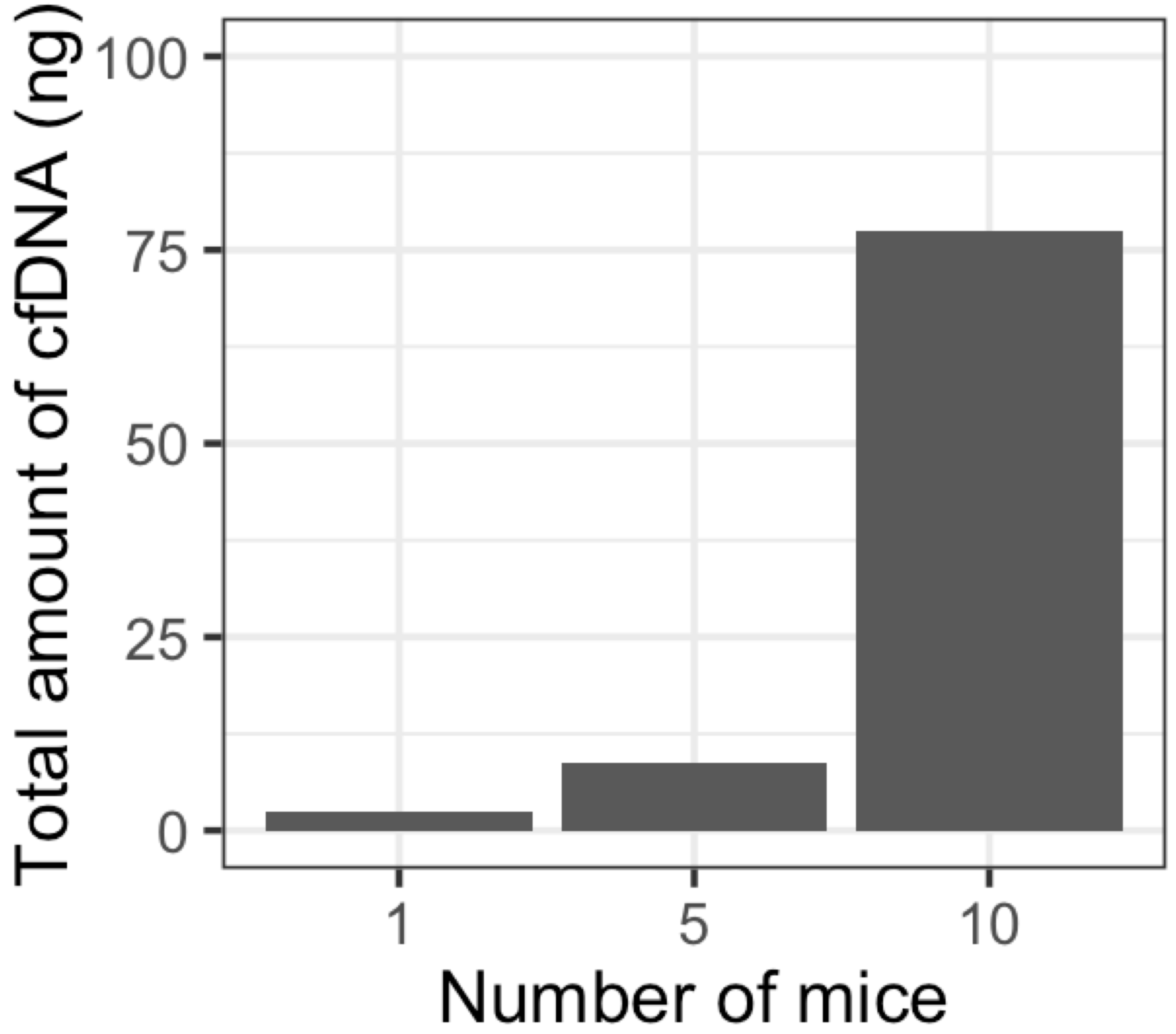

### Additional File 6

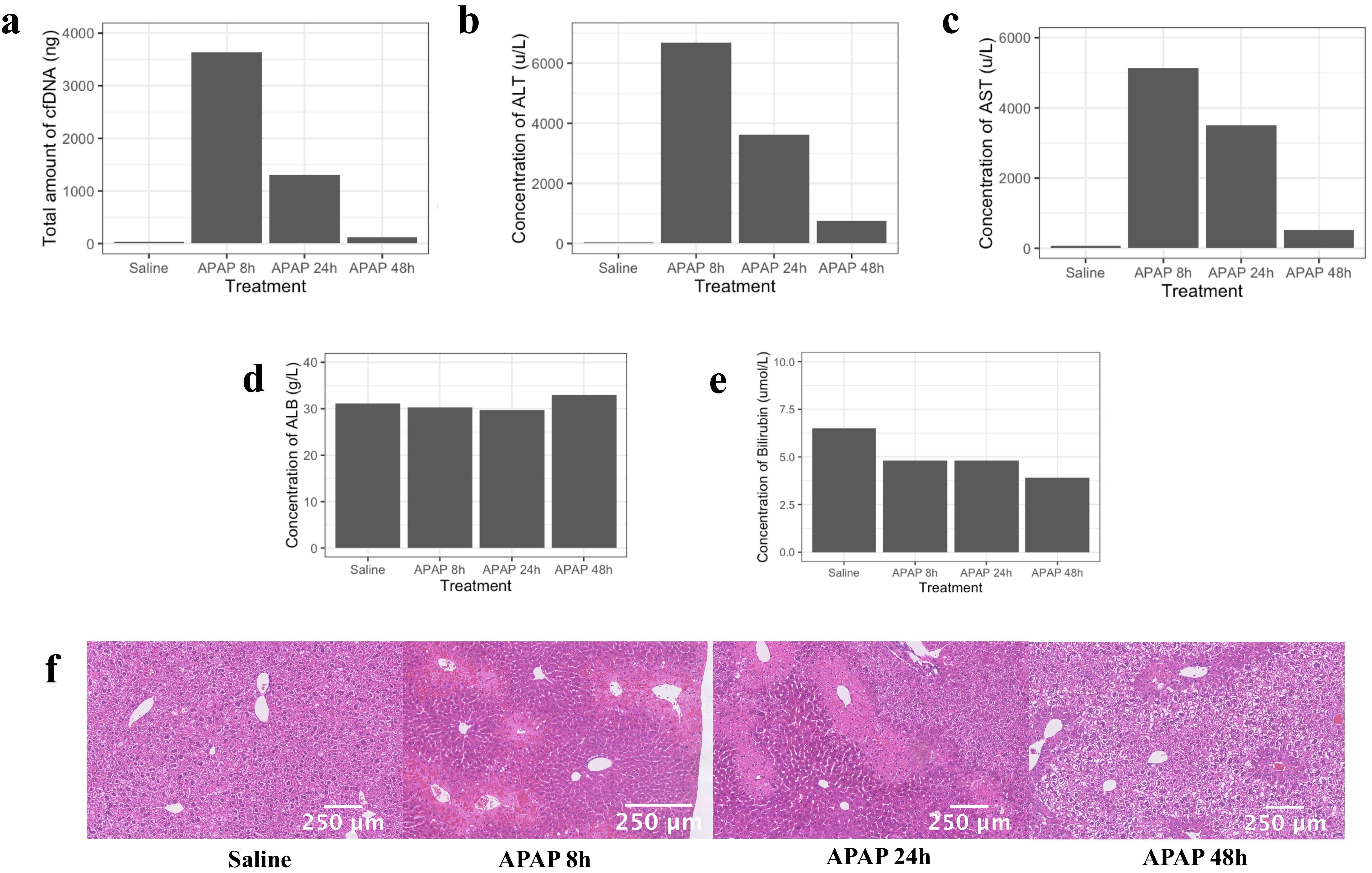

### Additional File 7

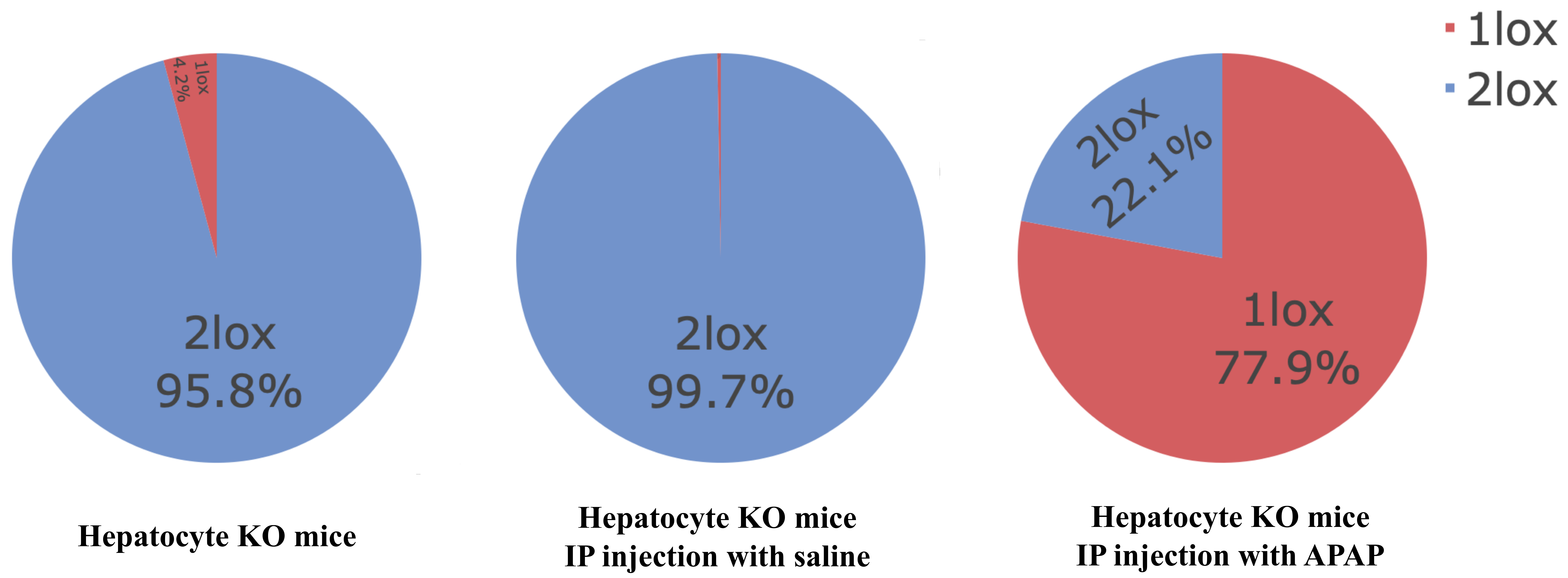
